# Reactivating a relaxation exercise during sleep to influence cortical hyperarousal in people with frequent nightmares – a randomized crossover trial

**DOI:** 10.1101/2025.03.11.642530

**Authors:** C. Sayk, A. Probst, F. Lange, S. Eickemeier, J. Amores, H.-V. Ngo, M. Ehsanifard, K. Junghanns, I. Wilhelm

**Affiliations:** Department of Psychiatry and Psychotherapy, Translational Psychiatry Unit, University of Luebeck, Luebeck, Germany; Institut of Psychology, University of Hildesheim, Hildesheim, Germany; Microsoft Research, Cambridge, MA, United States; Department of Psychology, University of Essex, Colchester, United Kingdom

**Author notes:** **Correspondence:** Ines Wilhelm Klinik für Psychiatrie und Psychotherapie Zentrum für Integrative Psychiatrie ZIP gGmbH | UKSH Campus Lübeck Universität zu Lübeck Ratzeburger Allee 160 23538 Lübeck.

## Abstract

**Study Objectives:** High-frequency EEG activity during sleep (cortical hyperarousal), is a transdiagnostic feature across psychiatric disorders, including nightmare disorder. It is discussed as a target of intervention; however, specific treatment options are yet unavailable. We tested whether exposure to relaxation-associated odor cues during sleep would reduce cortical hyperarousal, i.e. beta (16.25 – 31 Hz), gamma (31.25 – 45 Hz), spindle activity and nightmare occurrence in participants with frequent nightmares.

**Methods:** Twenty-five (21 female, mean age (SD) = 24.94(5.01)) participants, recruited from undergraduate students at University of Luebeck, with ≥1 nightmare / week received a deep breathing relaxation intervention for one week coupled with an odor. On two subsequent nights in the sleep laboratory, the associated odor (A), or control odor (B) were presented in randomized order in a crossover design with randomization at baseline; participants were blinded to intervention.

**Results:** N = 11 participants were allocated to AB and n = 14 to BA sequence. Exposure to relaxation-associated odor cues during sleep did not affect beta or gamma activity while spindle count and density were significantly reduced. Reduction in spindle count during reactivation nights correlated with reduced subjective wake-after-sleep-onset. There was no additional impact on nightmare symptoms. There were no adverse events or side effects.

**Conclusions:** The reactivation of relaxation-associated states with odor cues during sleep may be associated with changes in spectral activity, specifically spindle activity. Future studies should implement multiple nights of reactivation and include different patient groups with cortical hyperarousal to test the transdiagnostic potential of this new intervention.

## Introduction

Cortical hyperarousal can be defined as increased high-frequency electroencephalographic (EEG) activity during sleep, especially in the beta and gamma bands (Blaskovich et al., 2020). It occurs in multiple disorders such as insomnia (Zhao et al., 2021), posttraumatic stress disorder (PTSD) (Wang et al., 2020) and depression (Lin et al., 2023) and could therefore be considered a transdiagnostic feature of the abovementioned disorders. In addition, (cortical) hyperarousal is associated with symptom pathways and severity. Briere et al. (2015) found for example, that hyperarousal is predictive of suicidality in PTSD patients. Likewise, psychological hyperarousal symptoms, including sleep disturbances, are often residual symptoms even after successful PTSD treatment (Kovacevic et al., 2023; Larsen et al., 2019; Miles et al., 2022; Schnurr & Lunney, 2019) and can thus be considered as insufficiently targeted by standard treatment protocols.

Cortical hyperarousal has been implicated in etiology models of insomnia (Levenson et al., 2015; Riemann et al., 2015), nightmare disorder (Gieselmann et al., 2019) and PTSD (Krystal & Neumeier, 2009). In the latter it has been linked to increased locus coeruleus and noradrenergic activity which are considered a potential source for this type of spectral activity (Kelmendi et al., 2015). Van Someren (2021) proposed that cortical hyperarousal could be caused by fragmented rapid eye movement (REM) sleep, that fails to downregulate noradrenergic tone as would occur in healthy REM sleep to facilitate emotional processing (Goldstein & Walker, 2014). Both fragmented REM sleep and cortical hyperarousal are thought to be influenced by a mixture of genetic predisposition and early childhood adversity (Muscatello et al., 2020). However, most findings on cortical hyperarousal and psychiatric symptoms are correlational so far (e.g. Blaskovich et al., 2020) and it has been difficult to determine, whether it is the outcome of fragmented REM sleep, a byproduct or even the cause of fragmented REM sleep. It has been argued for example, that nocturnal cortical hyperarousal, which is closer to wake-like EEG activity, might cause higher responsiveness to external cues and thus disrupt sleep (Xu et al., 2022).

Taken together (cortical) hyperarousal is a relevant transdiagnostic feature that may not only be a relevant biological mechanism but also a target for novel interventions. However, interventions that specifically target cortical hyperarousal and could thus help us to better understand its role and develop new treatment options are not available yet.

There are several options for the manipulation of cortical hyperarousal that were considered for this study. Cortical stimulation, such as transcranial direct current stimulation (tDCS), can be used to influence high-frequency spectral activity (Kayarian, Jannati, Rotenberg & Santarnecchi, 2020). However, stimulation usually leads to an increase in the targeted frequency band which would be counterproductive to the desired direction of the intervention. Relaxation exercises such as deep breathing or progressive muscle relaxation can influence high-frequency EEG-activity (Kim, Rhee & Kang, 2014; Lee, Bhattacharya, Sohn & Verres, 2012) and thus have the potential to reduce cortical arousal. The effects of relaxation exercises on EEG-activity mostly pertain to effects observed during the actual exercise (Jacobs & Friedman, 2004) or to wake-EEG in the follow-up (e.g. Cheng et al., 2018). However, it is still unclear how these effects can be translated to target cortical hyperarousal during sleep. Targeted memory reactivation (TMR), (Oudiette & Paller, 2013) is a method that can be used to reactivate memory contents via an (often auditory or olfactory) cue during sleep that has been associated with the cue at encoding (Born & Wilhelm, 2012). Beck et al. (2021) demonstrated that exposure to relaxation-related words, such as “relax” or “sea,” during NREM sleep influenced spectral activity. More specifically, it reduced interhemispheric asymmetry of slow-wave activity (SWA) while enhancing overall slow-wave activity, suggesting that relaxation-associated concepts can be reactivated during sleep, thereby increasing sleep depth. Therefore, reactivating relaxation-associated content may also have the potential to influence cortical hyperarousal during sleep. Recent findings by Schwartz et al. (2022) found that targeted memory reactivation (TMR) of imagery rehearsal therapy (IRT) in individuals with frequent nightmares led to additional improvements in nightmare symptoms. However, this study did not report whether changes in spectral activity occurred in the TMR group. In sum, these findings suggest that TMR could be a feasible and beneficial approach for individuals experiencing frequent nightmares. However, these studies used auditory cues to reactivate memory contents. Auditory cues, in themselves, can influence spectral activity, leading to increased slow oscillatory and spindle activity (Ngo et al., 2013). This makes it challenging to determine whether observed effects on spectral activity are due to memory reactivation processes or simply a result of auditory stimulation. In contrast, olfactory cues do not lead to these changes in spectral activity. Moreover, unlike auditory cues, olfactory stimuli directly reach the hippocampus (Bar et al., 2020), increasing the likelihood of detecting any effects of TMR on arousal.

As mentioned above, cortical hyperarousal is increased in patients who suffer from frequent nightmares. More specifically, Blaskovich and colleagues (2020) found increased beta- and gamma activity, especially shortly before entering REM-sleep, in subjects with frequent nightmares. Additionally, individuals with frequent nightmares exhibited increased fast spindle activity compared to a control group (Picard-Deland et al., 2018). Recent findings from our lab corroborate the findings of Blaskovich et al. (2020). A group of nightmare patients, who received imagery rehearsal therapy (IRT) to treat nightmares (Thünker & Pietrowsky, 2021), showed increased cortical arousal compared to healthy controls, independent of actual nightmare occurrence during the measurement (Sayk, Saftien, Koch, Ngo, Junghanns & Wilhelm, 2024). Moreover, when comparing EEG-activity before and after the intervention, there was a trend towards lower gamma activity after imagery rehearsal therapy, which again points to the important role of cortical hyperarousal in nightmare aetiology. However, these findings are correlational and interventions that directly target cortical hyperarousal, such as a deep-breathing relaxation exercise, and include an experimental manipulation are still lacking. This is especially important given the transdiagnostic aspect of cortical hyperarousal and the potential of interventions that target it.

The primary goal of this exploratory study was to investigate whether reactivating a relaxation exercise in participants with frequent nightmares that has been practiced for one week led to 1) a reduction of cortical hyperarousal during sleep as indicated by beta, gamma and spindle activity and 2) a reduction in nightmare symptoms.

## Methods and Materials

### Participants

Participants were mainly recruited from the undergraduate student body at the University of Luebeck, Germany from 03/2022 to 01/2023 and ended when the pre-determined sample size was reached. For this experiment, we included individuals between 18 and 35 years old who have at least one nightmare a week, do not to suffer from any other psychiatric (especially affective disorders, anxiety or posttraumatic stress disorder) or somatic illness (esp. neurological disorders) or other sleep disorders (mainly insomnia, circadian rhythm disorders and sleep apnea) and do not take any sleep-altering medication. Eligibility criteria were determined in a telephone interview, with a special emphasis on ensuring that nightmare symptoms were present according to the following definition given to the potential participants: “Dreams with very negative emotional experiences which are usually so upsetting / frightening that you wake up from them”.

The study was approved by the university’s ethics committee. All participants provided written informed consent after they have been given a complete description of the study protocol. Participants were compensated financially (four participants received part of the compensation in the form of course credits) for study participation.

The initial sample consisted of 29 participants with frequent nightmares. Four participants dropped out of the study after adaptation night due to inability to sleep in the sleep lab. The majority of the initial sample (n = 21) was female, mean age was 23.58 (SD = 4.6). Participants mostly suffered from *nightmares since childhood* (n = 5) or for *at least 10 years* (n = 11), the rest reported an onset of nightmares for *at least five years* (n = 6) or for *one year or less* (n = 7). On a 5-point Likert scale, 20 participants rated their degree of suffering from nightmares as *“moderate”* and 5 participants reported a *“strong”* degree of suffering. Most of the participants remembered nightmare content very often or always. Six of the participants reported an identical or almost identical repetition of one nightmare, and nine participants reported their nightmares to relate to their biography. Participants on average had healthy sleep according to the Pittsburgh Sleep Quality Inventory (PSQI) and average sleep reactivity according to the Ford Insomnia Response to Stress Test (FIRST). Since suggestibility and absorption have been implicated in previous research on relaxation (Cordi & Rasch, 2020) we assessed all participants’ suggestibility, i.e. the tendency to react to hypnotic suggestions on the Harvard Group Scale of Hypnotic Susceptibility 5-Item Short-Version (HGSHS-5: G) and their absorption levels, i.e. imaginative involvement and the tendency to become mentally absorbed in everyday activities on the Tellegen Absorption Scale (TAS) (see also Table 1). Suggestibility levels in our sample can be rated as medium (Riegel et al., 2021) whereas levels of absorption were comparatively low (Angiulo & Kihlstrom, 1993). Sample size was determined using a medium to large effect size (cf. Cheng et al., 2018 for relaxation exercise; Cordi & Rasch, 2020 and Beck et al., 2021 for the effects of (reactivating) relaxation on sleep), 1-β = 0.85, α = 0.05 and correlation among repeated measures = 0.5, which resulted in N = 21 to which we added a 20% allowance for dropouts, resulting in a desired final sample size of N = 25.

**Table 1.**
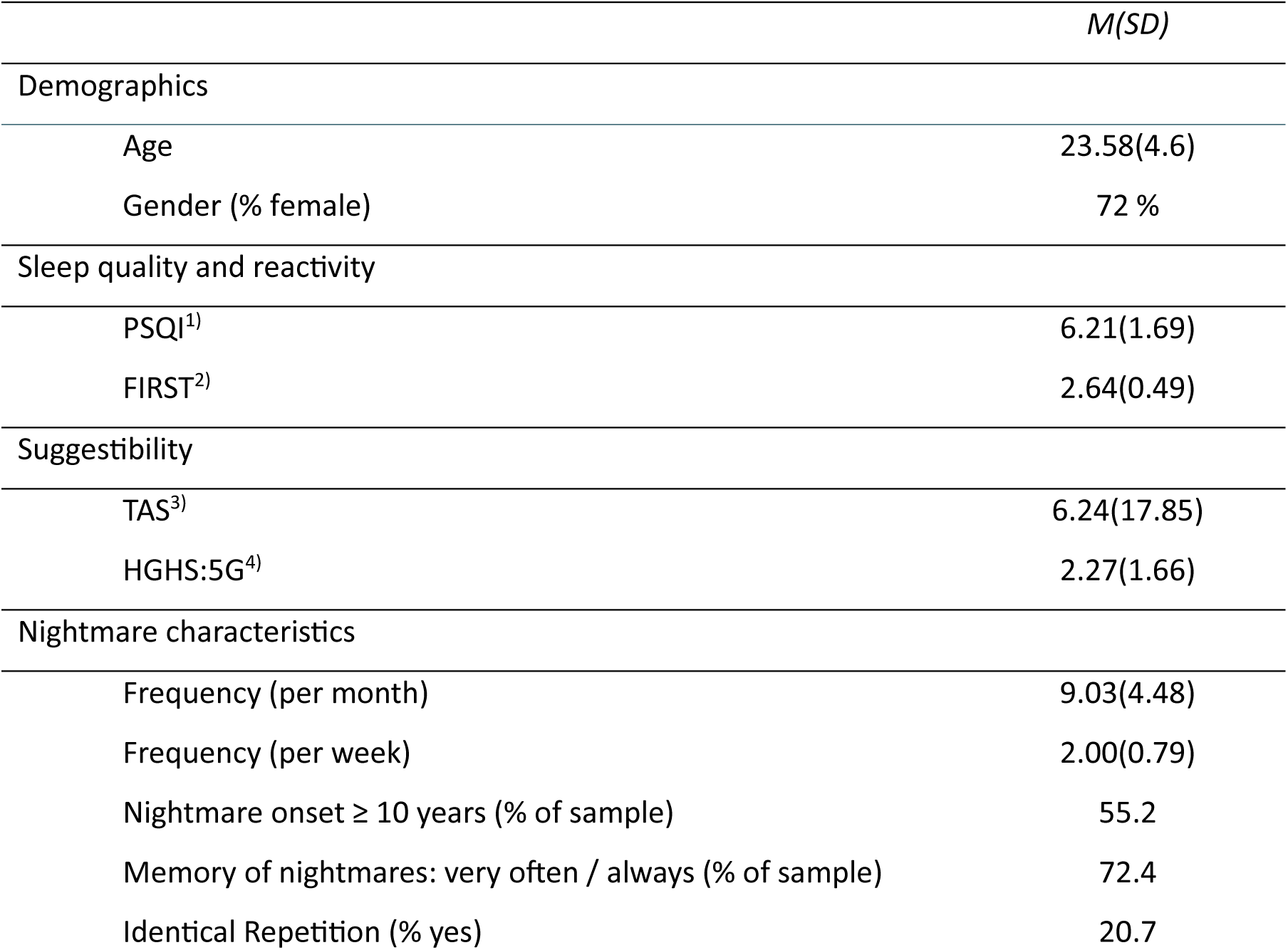
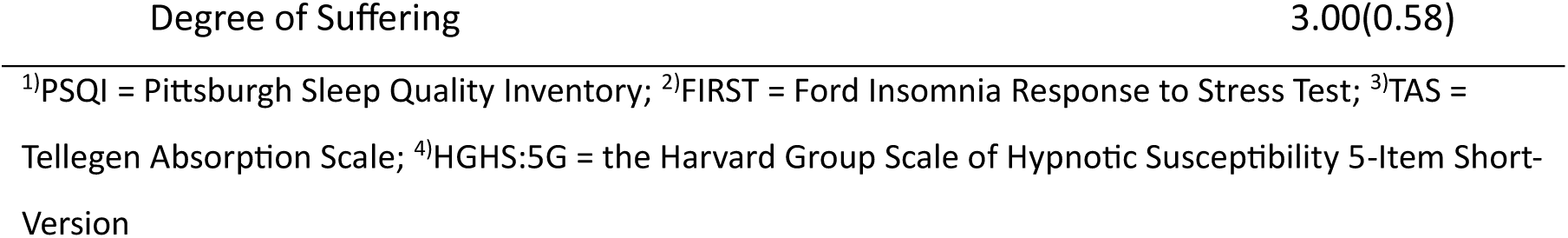
Demographic and psychometric characteristics of sample at baseline.

### Procedure and Design

#### Procedure

Potential participants were first screened for eligibility in a telephone interview, with a particular focus on ensuring the criterium of at least one genuine nightmare per week. If all inclusion and exclusion criteria were met, their three nights in the sleep laboratory, one adaptation night and two experimental nights, were scheduled (Figure 1). The first set of questionnaires were completed via an online link before coming to the sleep laboratory for their adaptation night. Upon arrival for the adaptation night, they rated four odors concerning pleasantness, familiarity, and intensity after having been presented with each odor directly from the container for 10 *s* each from a distance of 20 *cm*. The odors were plant-based scents (lemon grass, rose geranium, tea tree and frankinsense) but the labels were concealed from the participants. The two odors that were rated to be the least familiar but the most pleasant and intense were chosen as relaxation-associated and control odor respectively for that participant. In case of a conflict between familiarity and pleasantness, the more pleasant one was chosen to ensure participants’ compliance. Afterwards, they were fitted with EEG, EOG, EMG, ECG and SCR electrodes before completing the relaxation exercise and listening to a control text which were both followed by a 2 x 2 minutes block design of eyes open and eyes closed to assess the aftereffects of both audios. The audios were presented on a computer using PsychoPy 3.1 (Peirce, Hirst & McAskill, 2022) and this procedure was repeated on the evening of the first experimental night. Levels of relaxation as well as positive and negative affect were measured at baseline and after each audio. Relaxation was measured using a 5-point Likert scale, while positive and negative affect was evaluated using a short version of the Positive and Negative Affect Schedule (PANAS). The relaxation exercise and the control text were presented in randomized order across participants and nights (adaptation night and first experimental night). The computerized presentation of the audio recordings was used to precisely send triggers for each segment of the audios (two 2-minute blocks of eyes open and eyes closed) to the EEG recording. Then they went to bed at their usual bedtime and an adaptation odor was presented with a diffusor for the whole night. In the morning, participants downloaded the audio of the relaxation exercise on their phones and received a diffusor (see material section) filled with their individual relaxation-associated odor. They were instructed to practice the relaxation exercise daily for the next seven days, activating the diffusor each time. The relatively short seven-day interval, compared to two (Schwartz et al., 2022) or even three weeks (Pruiksma et al., 2016) in previous studies, was chosen to facilitate recruitment of participants and ensure continued compliance in this exploratory study design. At the end of this week of daily practice, participants completed another set of online questionnaires and returned to the sleep laboratory for the first of two experimental nights. On the evening of the first experimental night, they were exposed to the relaxation exercise and the control text. This time, the olfactometer (Amores, 2016) was activated when the participants first reached stable slow wave sleep. The order in which the odors were presented in the first and second experimental night was randomized between subjects and participants were blind to intervention. The last experimental night with odor presentation took place another 2 – 7 days after the first experimental night, depending on sleep laboratory capacities, so that participants could have at least one night of restorative sleep in their home sleep environment, but experimental nights were not too far apart either. (see Figure 1 for complete overview of the experimental procedure).

**Figure 1.**
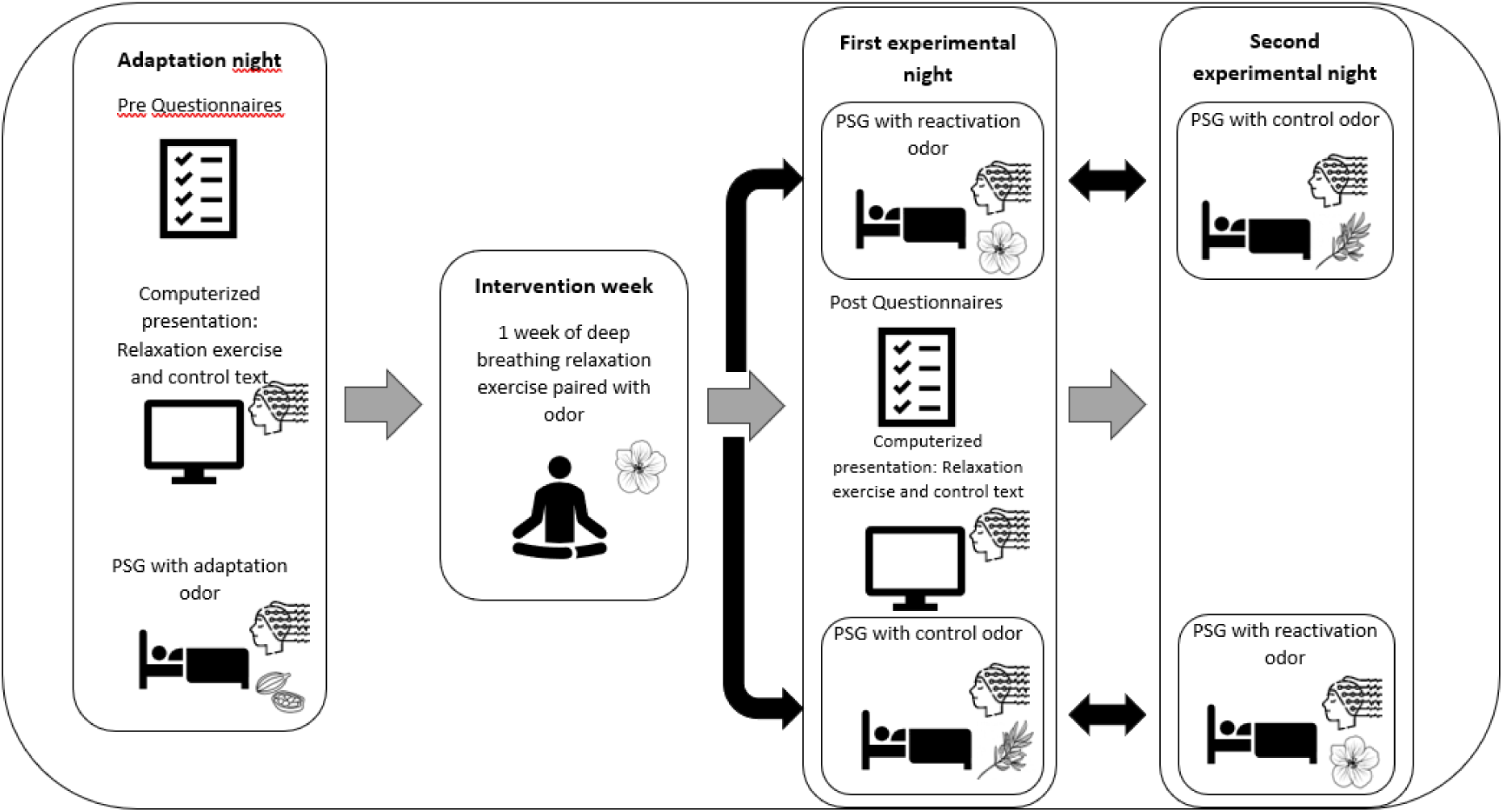
Experimental procedure. Schema of the study procedure and use of odors. At adaptation night, participants fill in the pre questionnaires, are presented relaxation exercise and control text at the computer in randomized order and a reactivation and a control odor are chosen according to participant’s rating. At adaptation night, a separate adaptation odor (cardamom) is presented to adapt participants to sleep while an odor is present. Participants then practice the deep breathing relaxation exercise at home for a week while activating the reactivation odor chosen for them (e.g. rose geranium). Then they return for two experimental nights where either the reactivation (e.g. rose geranium) or a control odor (e.g. tea tree) are presented in randomized order (possibilities for order of presentation are marked by black arrows). At the first experimental night, they also filled in post questionnaires and listen to the relaxation exercise and control text again.

#### Design

A crossover design was chosen to reduce interindividual variability, especially since spectral activity during sleep is known to have a high interindividual variance, even in healthy young adults (cf. Tucker et al., 2007). Odor presentation was delivered in an AB or BA sequence at the first and second experimental night (A = reactivation odor; B = control odor). Participants were randomly assigned to one of the two sequences at adaptation night according to randomly computer-generated numbers with n = 11 participants allocated to AB and n = 14 allocated to BA in the final sample. The computerized presentation of the audios for the relaxation exercise (A) and control text (B) at baseline (pre) and at the first experimental night (post) also followed a randomized order of presentation with the possible sequences AB (pre) – BA (post) and BA (pre) – AB (post), with n = 12 of the final sample allocated to AB – BA and n = 13 to BA – AB.

### Materials

#### Questionnaires

Nightmare symptoms were assessed with a customized questionnaire at baseline and after the intervention week including nightmare frequency, time of nightmare onset, nightmare content, ability to remember nightmares and distress caused by nightmares (see Sayk et al., 2024). Subjective sleep quality was measured with the *Schlaffragebogen A*, SF-A/R (Görtelmeyer, 2011) during sleep lab nights, and retrospectively using the Pittsburgh Sleep Quality Inventory, PSQI (Buyssee, Reynolds,

Monk, Berman & Kupfer, 1989) at baseline and after the intervention week. Additionally, at baseline, sleep reactivity was assessed with the Ford Insomnia Response to Stress Test (FIRST) (Drake et al., 2004) and suggestibility and absorption were measured with the Harvard Group Scale of Hypnotic Susceptibility 5-Item Short-Version (HGSHS-5: G) (Riegel et al., 2021) and Tellegen Absorption Scale (TAS) (Tellegen, 1992) respectively. Directly after the relaxation exercise and control text, participants repeatedly rated their current affective state with a short version of the Positive and Negative Affect Schedule (PANAS) (Breyer & Bluemke, 2016).

#### Polysomnography

Participants were fitted with 64 electrode EEG caps (ActiChamp, Brain Products, Gilching, Germany) referred to the left mastoid (M1) and ground electrode. We used electromyography (EMG) placed on the chin, as well as electrooculography (EOG) and electrocardiography (ECG) measurement. Data was recorded with Brain Vision Recorder (Brain Products, Gilching, Germany). Impedances were below 10 kΩ. Sampling rate was 500 Hz.

#### Sleep Scoring and spectral power analysis

Sleep stages were scored manually according to AASM criteria (Berry et al., 2020) by two experienced sleep lab technicians. Recordings were visually inspected on a 30-second basis and channels with muscle- and technical-related artifacts were discarded. Additionally, for the wake EEG data, eye movement artifacts were removed with independent component analysis (ICA). Artifact-free, 50% overlapping, 8.192-second epochs were Hanning-tapered and Fast Fourier Transformed (FFT) in order to calculate absolute power of spectral densities for each frequency bin between 1.25 Hz and 45 Hz for NREM (Stage 2 and Stage 3) and REM sleep periods, separately. Spectral power was calculated for wake EEG, the whole night, for odor-on and odor-off periods separately and for 10 minutes pre- and post-REM (cf. Blaskovich et al., 2020). Pre-REM periods were defined as 10-minute intervals of NREM sleep directly before the onset of the first two REM periods. Accordingly, post-REM periods included similar 10-minute NREM epochs following the end of the first two REM periods to ensure REM periods were comparable across all participants in timing and number and were followed by stable NREM sleep. Band-wise spectral power was extracted by summing up bin-wise values into ranges of beta (16.25–31 Hz) and gamma (31.25–45 Hz) bands for sleep EEG data and alpha (8.25–13 Hz), beta (16.25–31 Hz) and gamma (31.25–45 Hz) bands for wake EEG data gathered during the computerized presentation of the relaxation exercise and control text. Additionally, spindle count, density, amplitude and threshold were calculated for stage 2 and 3 of NREM sleep combined in the frequency band between 12 and 16 Hz. For exploratory purposes, delta (1.25 Hz – 4 Hz), theta (4.25–8 Hz), alpha (8.25– 13 Hz) and sigma (13.25–16 Hz) activity was analysed for sleep.

#### Olfactometer

During experimental nights, odors were presented using an olfactometer “Essence” (Amores, 2016), that was placed in a holder directed at the participant’s face at 50 *cm* distance. Odor presentation was started by the experimenter when the participants first reached slow wave sleep, to reduce the possibility of them waking up when odor presentation began and stopped by the experimenter 30 minutes before their scheduled wake-up time or when they showed signs of waking up before that on the PSG, thus encompassing both NREM and REM sleep. Odors were presented in 20 millisecond bursts every 55 seconds that were arranged in four-minute loops of odor-on and odor-off periods. These parameters were chosen so that potential changes in spectral activity associated with the odor bursts could be reliably detected and habituation effects were reduced via the intermittent odor presentation. For practice at home, participants received a commercially available diffusor.

#### Relaxation exercise and control text

The relaxation exercise used in this study was a custom-made 15-minute deep breathing exercise that enabled the participants to practice deep, diaphragmatic breathing and relax. On average, participants rated the ability to follow the instructions for the relaxation exercise to be high (*M* = 4.04, *SD* = 1.03) as well their ability to engage with the exercise at first trial (*M* = 4.01, *SD* = 0.97) on a 5-point Likert scale from 1 = “do not agree at all” to 5 = “completely agree”. For the week of relaxation exercise, 18 participants reported to have practiced daily, while those who did not practice daily missed 2.86 (SD = 2.03) days of practice on average. Mean ratings of understanding of and engagement with the exercise were 4.37 (SD = 0.71) and 4.16 (SD = 0.7) respectively.

As a control, participants listened to a text about polar research. This control text and the relaxation exercise were presented on a computer in the evening of the adaptation night and of the first experimental night. The order of the control text and the relaxation exercise was randomized across participants and time points. The audio recordings of the relaxation exercise and the control text were narrated by a member of the research team who was otherwise not involved in any interactions with the participants. Both recordings were delivered in a calm voice, carefully matched in tone and pacing. In the sleep laboratory, participants listened to the recordings through headphones at a fixed volume while seated with their eyes closed.

### Statistical analyses

Statistical analyses were carried out with SPSS and FieldTrip (Oostenveld, Fries, Maris & Schoffelen, 2011). Differences in sleep architecture were evaluated by dependent samples t-tests. Šidák correction was used to correct for multiple comparisons. Differences in band-wise spectral power for the computerized presentation of the relaxation exercise and control text, the whole night, odor-on vs. odor-off periods and 10 minutes pre- and post-REM (cf. Blaskovich et al., 2020) were tested with cluster-based permutation tests with an overall p-value set to p ≤ 0.05. The same tests were used for sleep spindle characteristics and correlation analyses between neural and behavioral measures. For the computerized presentation of relaxation exercise and control text, there were subsequent 2 x 2 condition (relaxation, control) and time of measurement (pre: adaptation night and post: first experimental night) analysis for the beginning (2^nd^ to 6^th^ minute) and end (8^th^ to 12^th^ minute). We chose to analyze EEG activity at the beginning and the end of the exercise separately based on previous studies demonstrating that EEG patterns during relaxation exercise differs between these two phases (Jacobs & Friedman, 2004; Lee et al., 2012). In line with these studies, we calculated mean EEG activity from the second to sixth minute defining the “beginning” and the final four minutes, excluding the last minute, defining the end (Jacobs & Friedman, 2004). Additionally, a 2(condition: relaxation and control) x 2 time of measurement (pre: adaptation night and post: first experimental night) x 2 blocks (eyes open, eyes closed) ANOVA was calculated to analyze the aftereffects of both audios. Changes in questionnaires before and after the intervention week were calculated with repeated-measure ANOVAs and correlation of questionnaire data and physiological data were calculated with cluster-based correlation analysis.

## Results

### Spectral activity during the relaxation exercise

When comparing spectral activity during listening to the relaxation exercise and the control text, a significant difference was observed at the beginning of the audios (2^nd^ to 6^th^ minute) with significantly less beta and gamma activity in the relaxation exercise than in the control text (figure 2). These effects did not persist to the end (8^th^ to 12^th^ minute) of the audios. There were no significant effects of time of measurement (pre: adaptation night and post: first experimental night) or any significant interactions between the factors condition (relaxation exercise and control text) and time of measurement. In the blocks of eyes opened and closed after the audios, a significant effect was observed for the alpha and beta band across both audios and time points with spectral activity being higher in the eyes closed compared to the eyes open. There were no significant effects of time of measurement (pre: adaptation night and post: first experimental night) or any significant interactions between the factors condition (relaxation exercise and control text) and time of measurement (pre: adaptation night and post: first experimental night).

**Figure 2.**
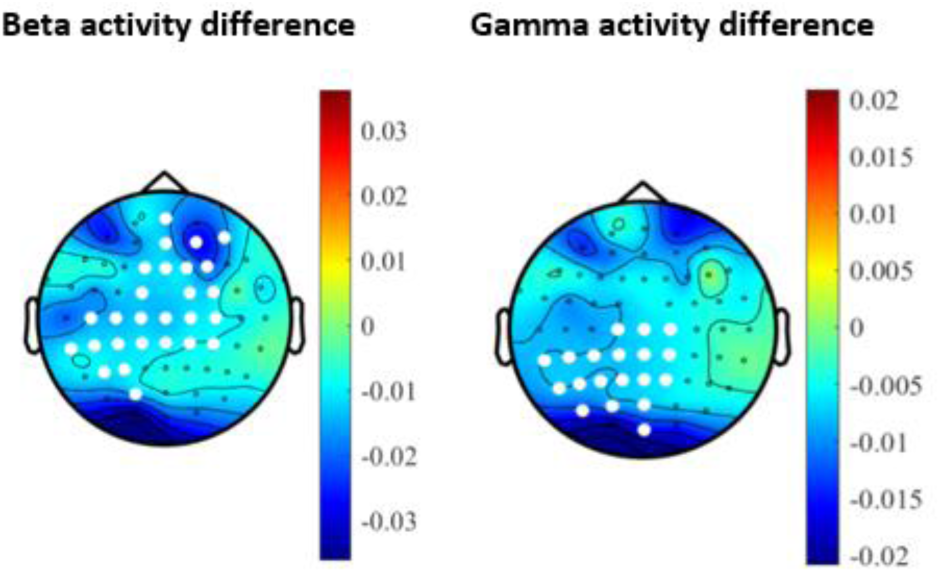
Difference in EEG activity between relaxation exercise and control text. Differences in beta (16.25 – 31 Hz, left) and gamma (31.25 – 45 Hz, right) activity in the beginning (2^nd^ to 6^th^ minute) of the relaxation exercise minus the beginning (2^nd^ to 6^th^ minute) of the control text. Significant electrode clusters (p < 0.05) are marked in white.

### Effects of relaxation exercise on subjective ratings

Subjectively rated relaxation rating relative to baseline relaxation ratings before listening to any of the audios was significantly higher after the relaxation exercise than after the control text (*F*(24,1) = 13.206, *p* = 0.001, η_p_² = 0.355; figure 3). There were no significant effects when comparing pre and post intervention week measurements (main effect of time of measurement) nor significant interactions between condition (relaxation exercise and control text) and time of measurement (pre: adaptation night and post: first experimental night). For the PANAS ratings, there was a significant effect of time on both positive and negative affect (*F*(21,1) = 18.479, *p* = 0.001, η_p_² = 0.468) with affect ratings generally being lower at the post than at the pre measurement. There was neither a significant effect of condition (relaxation exercise and control text) nor a significant interaction of condition (relaxation exercise and control text) x time of measurement (pre vs. post) on the PANAS ratings.

**Figure 3.**
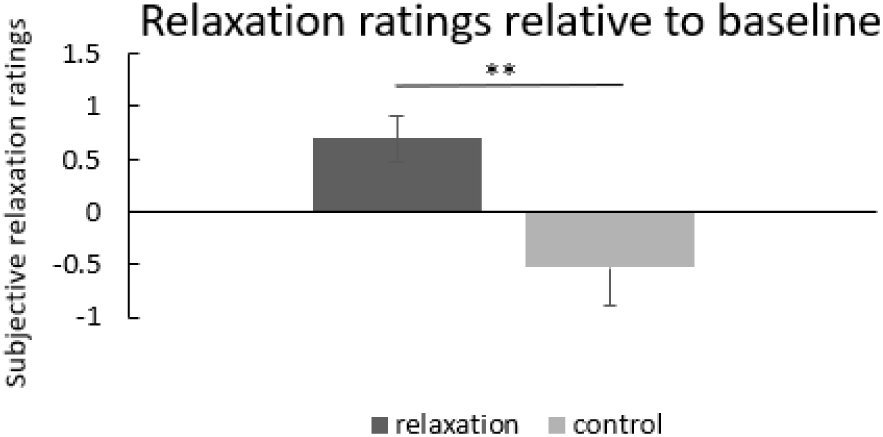
Relaxation ratings relative to baseline.

### Effects of reactivation on sleep architecture

The comparison of sleep architecture between nights with the reactivation odor and the control odor yielded no significant results (Table 2).

**Table 2.** Sleep architecture during nights with the reactivation and the control odor.

|  | Reactivation odor | Control odor |  |
| --- | --- | --- | --- |
|  | *M(SD)* | *M(SD)* | *p*-value |
| Total sleep time (minutes) | 435.46 (62.47) | 447 (61.7) | 0.285 |
| Sleep efficiency (%) | 91.35 (6.61) | 90.96 (9.64) | 0.776 |
| N1 duration (%) | 4.73 (2.42) | 5.09 (2.57) | 0.357 |
| N2 duration (%) | 52.02 (7.31) | 52.63 (8.96) | 0.622 |
| N3 duration (%) | 16.88 (5.29) | 17.46 (5.8) | 0.371 |
| REM duration (%) | 21.16 (3.85) | 20.75 (5.52) | 0.978 |
| REM density | 5.21 (3.16) | 4.23(2.15) | 0.126 |

Differences calculated by dependent samples t-test without the use of covariates, Šidák-adjusted $p$ -value = 0.0073.

### Effects of reactivation on spectral activity and spindle characteristics

When comparing spectral activity in reactivation and control nights for the entire night (for NREM and REM sleep separately), there were no significant differences in the beta and the gamma band. Comparing odor on and off periods within the reactivation nights did also not reveal any differences in the beta and gamma band neither during NREM nor REM sleep (Figure 4). Lastly, when examining spectral activity during the 10 minutes pre and post REM as has been done in previous studies (Blaskovich et al., 2020), there were neither significant differences for odor nor any significant interactions of odor with pre or post REM interval.

**Figure 4.**
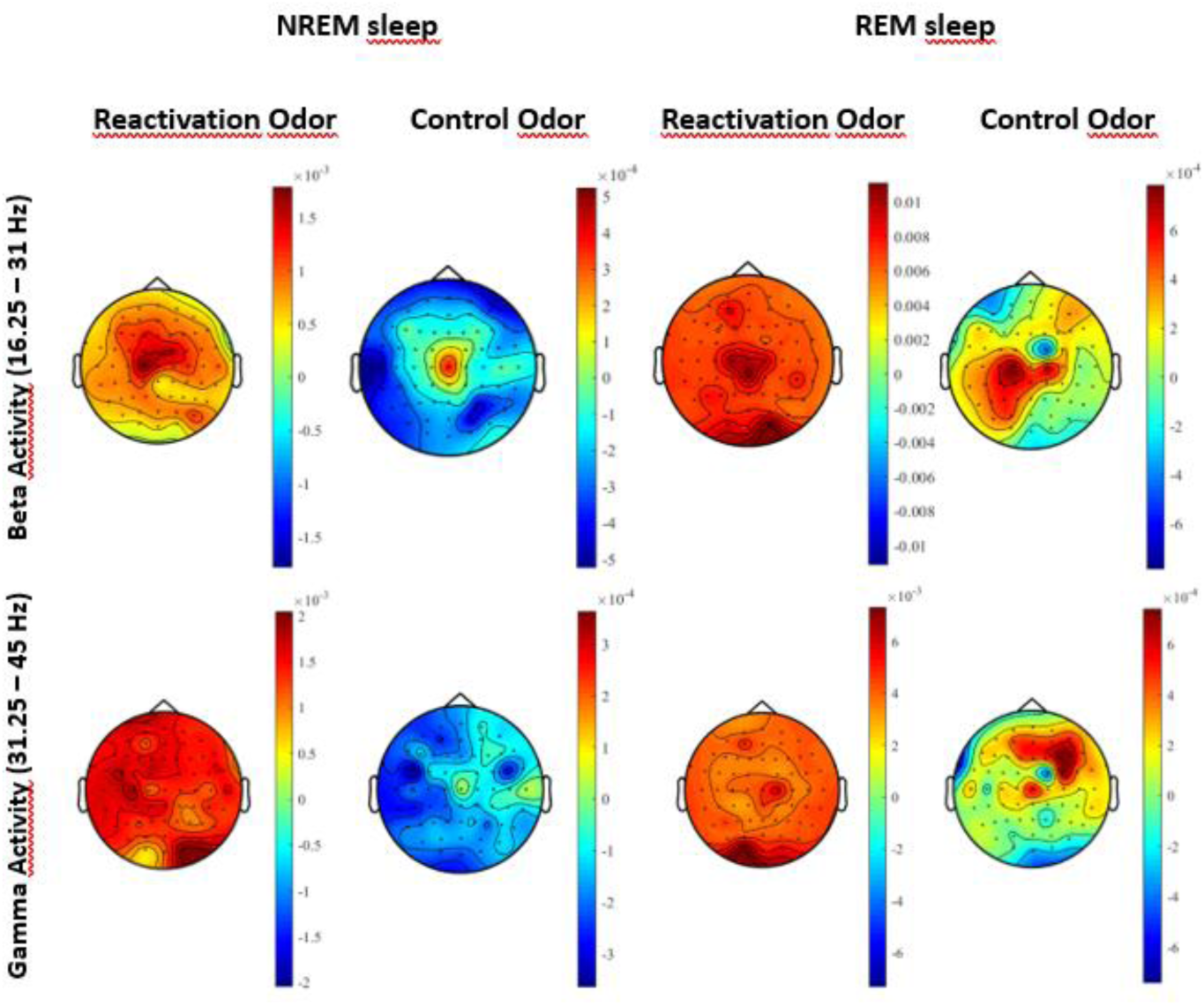
Beta and Gamma activity in NREM and REM sleep with and without reactivation. Absolute spectral power in the beta (16.25 – 31 Hz, top) and gamma band (31,25 – 45 Hz, bottom) with the reactivation odor on Absolute spectral power in the beta (16.25 – 31 Hz, top) and gamma band (31,25 – 45 Hz, bottom) did not differ between odor on and odor off periods neither for reactivation odor nor control odor.

*S*pindle count (Figure 5 C) and density (Figure 5 B) were significantly lower in reactivation nights than in control nights. There were no significant differences for spindle amplitude (Figure 5 A) and spindle threshold (Figure 5 D). There were no significant differences between odor on and odor off periods for spindle activity as analyzed with cluster-based statistics with a cluster-based significance level set to *p* ≤ 0.05.

**Figure 5.**
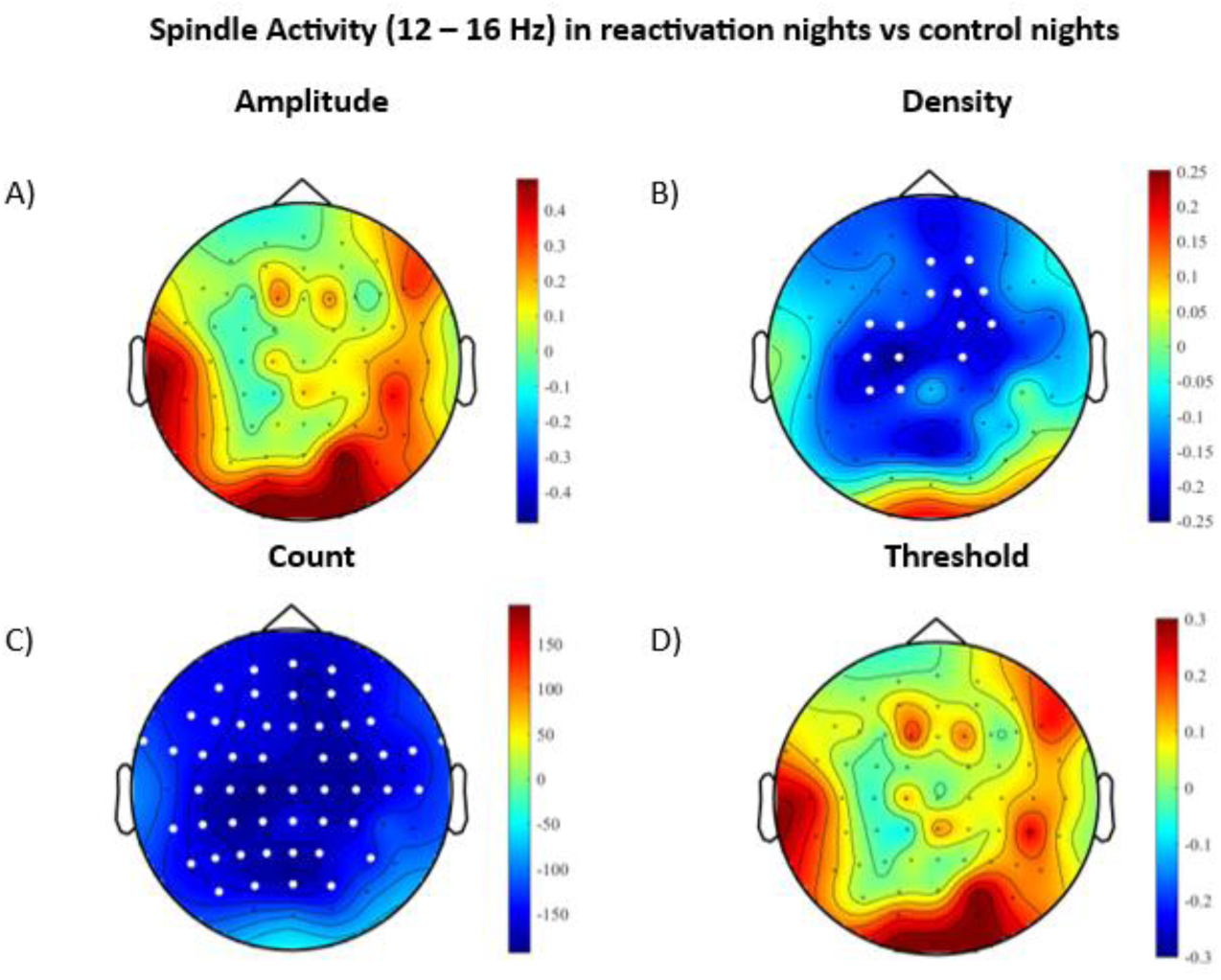
Spindle Activity (12 – 16 Hz) in reactivation nights vs control nights. Spindle parameters (12 – 16 Hz) between nights with the reactivation odor and nights with the control odor with significant electrode clusters marked in white (*p*-alpha < 0.05). Spindle count (C) and density (B) were lower when the reactivation odor was presented.

Exploratory analysis revealed that delta activity in NREM was significantly higher when the reactivation odor was present compared to when it was not, while this difference was not significant for the control odor. Moreover, there was no significant interaction between odor on and off periods and type of odor (relaxation odor vs. control odor). The observed effect was localized in a cluster of frontal and central electrodes (see Supplementary Material, Figure A). A similar pattern emerged in alpha activity during REM sleep, with alpha activity being higher during odor on phases of the reactivation odor compared to odor off phases. However, this difference was not significant for the control odor, nor was there a significant interaction between odor on and off periods and type of odor. The significant electrode cluster for this effect was located in a parieto-occipital cluster of electrodes (see Supplementary Material, Figure B). No other significant effects were observed for any other comparison or frequency band during sleep.

### Effects of relaxation exercise on nightmare symptoms and sleep quality

One week of practicing the relaxation exercise at home did neither affect subjectively rated retrospective sleep quality as indicated by the PSQI nor nightmare frequency or the level of distress caused by nightmares (all *p*s ≥ 0.417 for the comparison of the measurement before and after the week of practice; Table 3).

In addition to this retrospective assessment of nightmare characteristics and sleep quality during one week of relaxation exercise, we investigated the direct effect of reactivation in the sleep lab on these parameters. A comparison of nightmare occurrence, sleep quality, and both negative and positive affect between nights with the reactivation odor and those with the control odor revealed no significant differences. (all *p*s ≥ 0.249) (Table 4).

**Table 4.** Comparison between nights with reactivation and control odor.

|  | Reactivation odor | Control odor |  |
| --- | --- | --- | --- |
|  | *M(SD)* | *M(SD)* | *p*-value |
| Subjective sleep quality (SF-A/R)<sup>1)</sup> |  |  |  |
| sleep quality | 2.59 (0.66) | 2.47 (0.69) | 0.484 |
| trouble falling asleep | 3.68 (0.98) | 3.72 (0.98) | 0.870 |
| trouble staying asleep | 3.24 (1.25) | 3.5 (1.11) | 0.249 |
| sleep depth | 3.47 (0.58) | 3.32 (0.84) | 0.460 |
| relaxation after sleep | 3.38 (0.45) | 3.31 (0.67) | 0.642 |
| Nightmare occurrence (%) | 25 | 41 | 0.459 |

<sup>1)</sup>SF-A/R = *Schlaffragebogen A*; Differences in subjective sleep quality calculated by dependent samples t-test without the use of covariates, Šidák-adjusted $p$ -value = 0.01; Differences in nightmare occurrence calculated by Chi<sup>2</sup>-test

### Correlations between neural and behavioral data

For all significant differences on neural measures (i.e. beta and gamma activity during the relaxation exercise and spindle count and density during reactivation in sleep), we tested whether they correlated with any of the related behavioral data, which consisted of subjective relaxation and affect ratings during the relaxation exercise and subjective sleep quality for reactivation and control nights. For the relaxation exercise, there were no significant correlations between beta and gamma activity and behavioral data (i.e. subjective relaxation ratings and PANAS ratings) all *p*s ≥ 0.05 for the cluster-based permutation test statistics. However, spindle count across all nights, i.e. both relaxation and control nights combined, was positively correlated with trouble staying asleep so that a higher spindle count was associated with longer subjectively reported wake after sleep onset (Figure 6).

**Figure 6.**
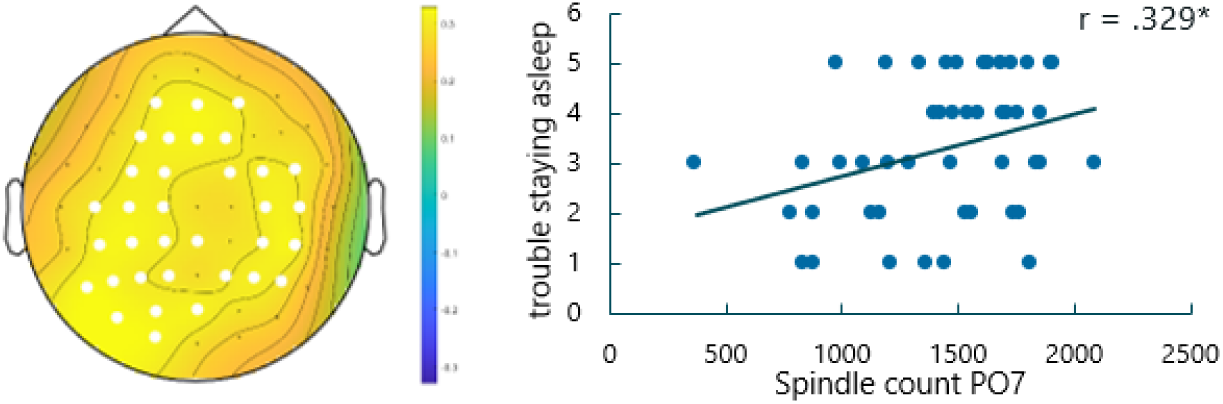
Correlation of spindle count and trouble staying asleep. Correlation of subjectively rated wake after sleep onset and spindle count with scatterplot for PO7 electrode. Significant electrode clusters are marked in white (*p*-alpha < 0.05).

### Harms and unintended effects

Apart from four participants dropping out due to sleep quality issues after the adaptation night in the sleep laboratory, there were no harms or unintended effects in this study.

## Discussion

The main goal of this randomized controlled exploratory study was to determine whether a week of relaxation exercise, which was reactivated during sleep using odors, reduces cortical hyperarousal indicated by beta and gamma activity, has an impact on spindle activity and reduces nightmare symptoms. While the relaxation exercise led to reduced beta and gamma activity during the exercise, this effect did not directly translate to the reactivation of the relaxation exercise, i.e. there was no effect of the reactivation on beta and gamma activity during sleep. However, the reactivation led to reduced spindle count and density, with spindle count being positively correlated with subjectively rated trouble staying asleep. No further effects on nightmare symptoms could be observed.

Consistent with previous studies (Cheng et al., 2018), the effects of the relaxation exercise on cortical hyperarousal and subjective ratings of relaxation while the participants were performing the exercise demonstrate that the deep breathing exercise functioned as expected. However, retrospective ratings of sleep quality and nightmares did not improve after practicing the relaxation exercise. Pruiksma and colleagues (2016) could show that a treatment focused on reducing arousal, which included relaxation exercises, reduced nightmare frequency and improved sleep quality in a way that was not inferior to a treatment arm including exposure and rescripting. However, studies that attempt to disentangle/decipher the effective components of nightmare treatments in general and the role of relaxation exercises in particular are still missing (Gieselmann et al. 2019). Therefore, it is difficult to determine the effects of the deep breathing exercise in the dosage used in this study. One major difference to the study by Pruiksma et al. (2016) is indeed dosage. Their intervention lasted three weeks compared to just one week which could explain the lack of effects on nightmare symptoms and sleep quality in our study. On a more conceptual level, it could also be possible that the relationship between cortical hyperarousal and nightmare symptoms is not causal and therefore altering spectral activity cannot alter nightmare symptoms. However, additional research with improved study designs would be needed to be able to draw a conclusion on that matter.

While a reduction in beta and gamma activity was observed during the relaxation exercise, reactivating the relaxation exercise during the night did not lead to a corresponding reduction in these frequency bands. One possible explanation is that baseline beta and gamma activity, i.e. in the control night, in our sample may have been lower than in previous studies, potentially due to a lower severity of nightmare symptoms. For example, the degree of suffering from nightmares prior to intervention in our current sample was *M(SD)* = 3.00 (0.57) while it was *M(SD)* = 4.04 (0.57) in our previous study where changes in gamma activity were detected following a nightmare-targeting intervention (Sayk et al., 2024). As a result, a further reduction due to the intervention may have been more difficult to detect in our sample. As the approach of this study is still very new and there is very little research on altering beta and gamma activity during sleep, it might also be that our approach was not well-suited to directly target these frequency bands and effects are instead displayed in spindle activity, which is discussed in more detail below. Exploratory analyses revealed that delta activity during NREM and alpha activity during REM were elevated when the reactivation odor was present. The increase in delta activity may be linked to increased relaxation, similar to the findings of Beck et al. (2021) who reported increased slow-wave sleep after presenting relaxation-associated cues during sleep. Elevated delta activity could therefore indicate deeper sleep and serve as an indirect marker of reduced arousal. Similarly, increased alpha activity in REM sleep during odor presentation may also reflect enhanced relaxation, as previous research has associated increased alpha activity during relaxation exercises with a state of greater relaxation (cf. Bing-Canar et al., 2016). However, these results should be interpreted with caution as the interaction of odor on and odor off phases and type of odor (relaxation odor vs. control odor) was not significant. This suggests that the observed effect may not entirely specific to the process of reactivation.

Beta and gamma activity were not influenced by the reactivation, spindle count and density were, however, reduced in reactivation nights. At first glance, these results appear to be unexpected, as they do not align with findings from previous reactivation studies, some of which report an increase in spindle activity in response to reactivation (Ngo et al., 2015). However, these studies differ significantly from ours in that they reactivated semantic contents, such as learned words, in healthy controls, rather than the content of a relaxation exercise in individuals suffering from frequent nightmares. Our results fit well within the emerging picture provided by findings on the relationship between spindle activity, arousal, and mental disorders characterized by heightened arousal. For example, Picard-Deland and colleagues (2018) found elevated fast spindle activity in subjects with frequent nightmares compared to healthy controls, and linked altered spindle activity to dream content, fear during dreams as well as anxiety and depression. When broadening the picture, similar findings have been presented for posttraumatic stress disorder in a review by Natraj and Richards (2023), that linked increased fast spindle activity to PTSD symptoms and suggested this increase to be caused by an increase in arousal. This notion is in line with results of Wang, Laxminarayan and colleagues, who found higher slow- and fast-spindle frequencies in veterans with PTSD compared to veterans with no PTSD symptoms (2020). They propose this increased spindle activity could be an expression of unsuccessful over-consolidation of a traumatic event. It is therefore possible that a similar maladaptive over-consolidation may occur in the context of nightmare disorders (van der Heijden et al., 2022).

Taken together, it seems likely that a) increased spindle activity is a nightmare related form of cortical hyperarousal that could have been present in our study sample as well and b) that this form of cortical hyperarousal was indeed influenced by the reactivation of the relaxation exercise. The positive correlation between spindle count and troubles staying asleep could suggest that a breathing exercise not only reduces spindle activity but also potentially improves subjective sleep quality.

Aside from our findings suggesting a correlational link between spindle activity and subjective sleep quality, we found no general effects of reactivation on any behavioral measures during reactivation in the sleep lab. More specifically, neither nightmare symptoms nor subjective or objective sleep quality was directly affected by the reactivation. Several factors might account for these lacking effects. First, the only time the reactivation occurred was in the sleep laboratory environment and research has shown that nightmare experience in the sleep lab is less severe and emotional than in the home sleep environment (Paul et al., 2019). Therefore, it could be advisable to measure the effects of reactivation for more than one night and outside a sleep laboratory in future studies. Other than the laboratory environment, the amount of reactivation could have been an issue, that should especially be considered when trying to expand this type of intervention to other disorders than nightmare disorder that involve cortical hyperarousal. Since we only reactivated the relaxation exercise for one night, this might not have been a long enough amount of reactivation. This becomes more apparent when taking into account other studies that reactivated psychotherapeutic interventions, as the successful ones tend to have longer periods of reactivation. For instance, a recent study by Schwartz and colleagues (2022) who reactivated imagery rehearsal therapy (IRT) by using auditory cues for a period of two weeks found significant effects of the reactivation, whereas Rihm et al. (2016) who reactivated exposure therapy for specific phobias using an odor cue for one night did not find additional effects on anxiety symptoms. Interestingly, they did, however, report changes in spectral activity after the reactivation which, taken together with our findings, might suggest that changes in spectral activity could precede behavioral or symptom changes in a reactivation context.

## Limitations

Mainly due to the exploratory and novel approach of this study, there are several limitations. In order to not overburden participants, we chose only one week of relaxation practice coupled with the odor. However, this may have resulted in a weaker association between relaxation and odor and potentially not enough practice of the deep breathing relaxation exercise (cf. Pruiksma et al., 2016) which both might have resulted in the incomplete effects on spectral activity and nightmare symptoms. In a similar vein and as discussed above, the reactivation for only one night, which was beneficial for the within-subjects study design, might have not been sufficient to influence nightmare symptoms (cf. Schwartz et al., 2022). Another potential issue in our study is that the washout period of 2–7 days may have been too short for some participants. This could explain why the effects of odor on delta and alpha activity were not strictly confined to the presence of the reactivation odor.

Future studies should therefore consider longer reactivation periods, possibly with portable olfactory devices. Mobile devices allow for targeted reactivation in the home-sleep environment, where nightmare symptoms might be more pronounced (Paul et al., 2019) thus potentially yielding larger intervention effects and allowing for a more direct transfer to therapeutic applications. Moreover, timing and type of intervention could be optimized so that the intervention targets both cortical hyperarousal and has the potential to influence specific symptoms, such as IRT for nightmares. Due to its transdiagnostic nature, TMR-based interventions that target cortical hyperarousal should be trialed in other disorders next, such as insomnia, PTSD or anxiety disorders. Lastly, one other route for future research could be drawn from Paul and colleagues (2019), that found changes in autonomic arousal directly related to nightmare occurrence. If a stable pattern of nightmare specific autonomic arousal was found, reactivation could be made even more targeted with the online detection of nightmare biomarkers, mainly in REM sleep, and subsequent direct reactivation of an effective intervention.

## Conclusion

The reactivation of a deep breathing relaxation exercise in participants with frequent nightmares did influence spindle parameters but not nightmare symptoms. However, the decrease in spindle count and density suggests that a partial increase in relaxation and decrease in nightmare-related neuronal activity occurred. As these mixed results could have been caused by type of intervention and length of the reactivation, future studies should explore longer reactivation periods and different intervention options to influence cortical hyperarousal in multiple disorders.

## Data availability statement

The data underlying this article will be shared on reasonable request to the corresponding author.

