## Supplementary Figures A and B for "Reactivating a relaxation exercise during sleep to influence cortical hyperarousal in people with frequent nightmares – a randomized crossover trial"

### Supplementary material

|  | **Delta Activity in NREM (1.25 – 4 Hz)** | |
| --- | --- | --- |
|  | **Reactivation Odor** | **Control Odor** |
| **Odor Presentation Period (ON vs. OFF)** | 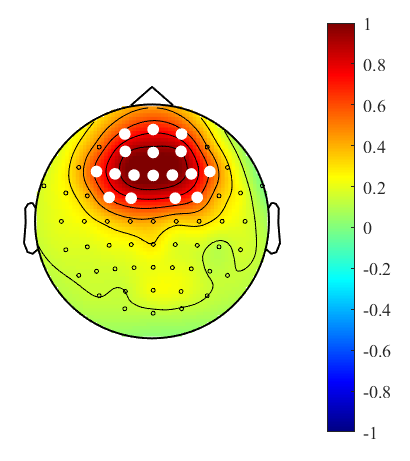 | 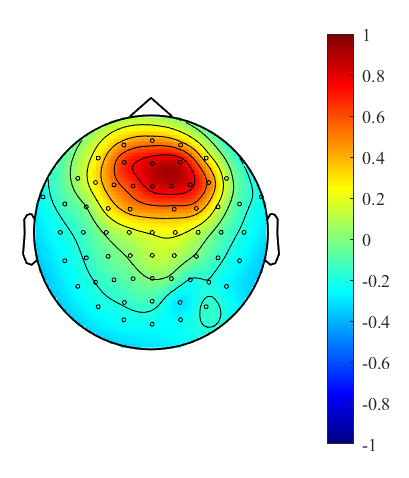 |
| **Figure A**. *Delta activity in NREM sleep with and without reactivation odor.*  Spectral Power differences in the delta band (1.25 – 4 Hz) between odor ON and odor OFF periods in the reactivation odor and the control odor with significant electrode clusters marked in white (*p*-alpha < 0.05). Delta activity was higher when the reactivation odor was presented. | | |

|  | **Alpha Activity in REM (8.25 – 13 Hz)** | |
| --- | --- | --- |
|  | **Reactivation Odor** | **Control Odor** |
| **Odor Presentation Period (ON vs. OFF)** | 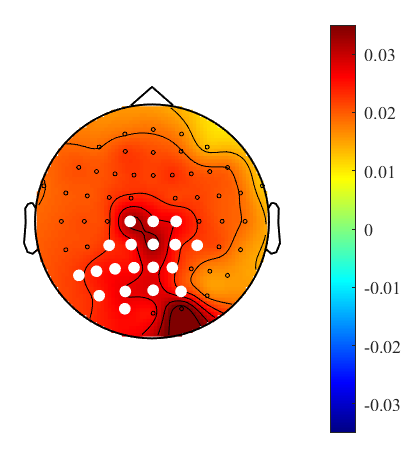 | 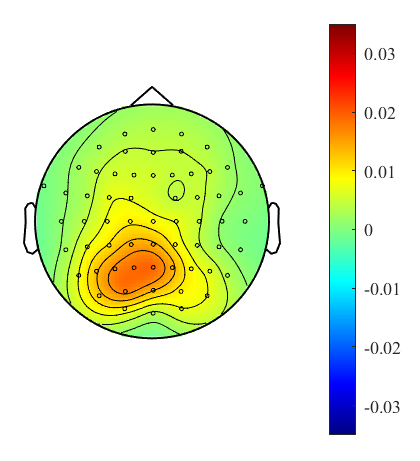 |
| **Figure B**. *Alpha activity in REM with and without reactivation odor*  Spectral Power differences in the alpha band (8.25 – 13 Hz) between odor ON and odor OFF periods in the reactivation odor and the control odor with significant electrode clusters marked in white (*p*-alpha < 0.05). Alpha activity was higher when the reactivation odor was presented. | | |
